# Mutations In *PIK3C2A* Cause Syndromic Short Stature, Skeletal Abnormalities, and Cataracts Associated With Ciliary Dysfunction

**DOI:** 10.1101/488411

**Authors:** Dov Tiosano, Hagit Baris Feldman, Anlu Chen, Marrit M. Hitzert, Markus Schueler, Federico Gulluni, Antje Wiesener, Antonio Bergua, Adi Mory, Brett Copeland, Joseph G. Gleeson, Patrick Rump, Hester van Meer, Deborah A. Sival, Volker Haucke, Josh Kriwinsky, Karl X. Knaup, André Reis, Nadine N. Hauer, Emilio Hirsch, Ronald Roepman, Rolph Pfundt, Christian T. Thiel, Michael S. Wiesener, Mariam G. Aslanyan, David A. Buchner

## Abstract

*PIK3C2A* is a class II member of the phosphoinositide 3-kinase (PI3K) family that catalyzes the phosphorylation of phosphatidylinositol (PI) into PI(3)P and the phosphorylation of PI(4)P into PI(3,4)P2. We identified homozygous loss-of-function mutations in *PIK3C2A* in children from three independent consanguineous families with short stature, coarse facial features, cataracts with secondary glaucoma, multiple skeletal abnormalities, neurological manifestations, among other findings. Cellular studies of patient-derived fibroblasts found that they lacked PIK3C2A protein, had impaired cilia formation and function, and demonstrated reduced proliferative capacity. Collectively, the genetic and molecular data implicate mutations in *PIK3C2A* in a new Mendelian disorder of PI metabolism, thereby shedding light on the critical role of a class II PI3K in growth, vision, skeletal formation and neurological development. This discovery expands what is known about disorders of PI metabolism and helps unravel the role of *PIK3C2A* and class II PI3Ks in health and disease.

## Introduction

Identifying the genetic basis of diseases with Mendelian inheritance provides insight into gene function, susceptibility to disease, and can guide the development of new therapeutics. To date, ∼50% of the genes underlying Mendelian phenotypes have yet to be discovered (Chong et al., 2015). The disease genes that have been identified thus far have led to a better understanding of the pathophysiological pathways and to the development of medicinal products approved for the clinical treatment of such rare disorders (Boycott et al., 2013). Furthermore, technological advances in DNA sequencing have facilitated the identification of novel genetic mutations that result in rare Mendelian disorders (Koboldt et al., 2013; Sobreira et al., 2015). We have applied these next-generation sequencing technologies to discover mutations in *PIK3C2A* that cause a newly identified genetic syndrome consisting of dysmorphic features, short stature, cataracts and skeletal abnormalities.

*PIK3C2A* is a class II member of the phosphoinositide 3-kinase (PI3K) family of lipid kinases that catalyzes the phosphorylation of phosphatidylinositol (PI) (Cantley, 2002). The functions of class II PI3Ks are poorly understood. However, they are generally thought to catalyze the phosphorylation of PI and/or PI 4-phosphate [PI(4)P] to generate PI(3)P and PI(3,4)P2, respectively (Jean and Kiger, 2014). PIK3C2A has been attributed a wide-range of biological functions including glucose transport, angiogenesis, Akt activation, endosomal trafficking, phagosome maturation, mitotic spindle organization, exocytosis, and autophagy (Behrends et al., 2010; Campa et al., 2015; Devereaux et al., 2013; Falasca and Maffucci, 2012; Franco et al., 2014; Gulluni et al., 2017; Krag et al., 2010; Leibiger et al., 2010; Posor et al., 2013; Yoshioka et al., 2012). In addition, PIK3C2A is critical for the formation and function of primary cilia (Falasca and Maffucci, 2012; Franco et al., 2014). However, there is as yet no causal link between *PIK3C2A*, or any class II PI3K, and human disease. Here, we describe the evidence that homozygous loss-of-function mutations in *PIK3C2A* cause a novel syndromic disorder involving neurological, visual, skeletal, growth, and occasionally hearing impairments.

## Results

Five individuals between the ages of 8 and 21 from three unrelated consanguineous families were found by diagnostic analyses to have a similar constellation of clinical features including dysmorphic facial features, short stature, skeletal and neurological abnormalities, and cataracts (Figure 1, Table 1, Table S1). The dysmorphic facial features included coarse facies, low hairline, epicanthal folds, flat and broad nasal bridges, and retrognathia (Figure 1B, Table S1). Skeletal findings included scoliosis, delayed bone age, diminished ossification of femoral heads, cervical lordosis, shortened fifth digits with mild metaphyseal dysplasia and clinodactyly, as well as dental findings such as broad maxilla incisors, narrow mandible teeth, and enamel defects (Figures 1C, 1D, Table S1, Figure S1). Most of the affected individuals exhibited neurological involvement including developmental delay and stroke. This was first seen in individual I-II-2 when she recently started having seizures, with an EEG demonstrating sharp waves in the central areas of the right hemisphere and short sporadic generalized epileptic seizures. Her brain MRI showed a previous stroke in the right corpus striatum (Figure 1F). Hematological studies were normal for hypercoagulability and platelet function (Table S2). In addition, brain MRI of patient II-II-3 showed multiple small frontal and periventricular lacunar infarcts (Figure S1E). Unclear episodes of syncope also led to neurological investigations including EEG in individual III-II-2, without any signs of epilepsy. Her brain MRI showed symmetrical structures and normal cerebrospinal fluid spaces but pronounced lesions of the white matter (Figure S1E). Other recurrent features included hearing loss, secondary glaucoma, and nephrocalcinosis.

**Table 1.** Phenotypic characteristics of *PIK3C2A* deficient patients.

|  |  |  |  |  |  |
| --- | --- | --- | --- | --- | --- |
| Family | I | I | II | II | III |
| Patient | II-1 | II-2 | II-2 | II-3 | II-2 |
| Age (years) | 13 | 8 | 12 | 10 | 20 |
| Gender | female | female | male | male | female |
| Origin | Israel<br>(Muslim-Arabic) | Israel<br>(Muslim-Arabic) | Syria | Syria | Tunisia |
| Consanguineous | + | + | + | + | + |
| Height | -3.3 SD | -2.3 SD | -2.5 SD | -4.8 SD | -2.5 SD |
| Weight | -0.2 SD | -1.7 SD | -0.2 SD | -3.9 SD | -1.9 SD |
| Head circumference | -0.25 SD | N.D. | +0.9 SD | -1.1 SD | +0.5 SD |
| Congenital cataract | + | + | + | + | + |
| Secondary glaucoma | + | + | + | + | - |
| Hearing loss | + | - | - | + | + |
| Scoliosis/Skeletal abnormalities | + | + | + | + | + |
| Teeth Abnormalities | + | + | + | + | + |
| Developmental delay | + | + | N.D. | N.D. | N.D. |
| Elevated urine MPS levels | + | N.D. | + | + | N.D. |

"+" indicates presence of trait, "-" indicates absence of trait, N.D., not done.

**Figure 1.**
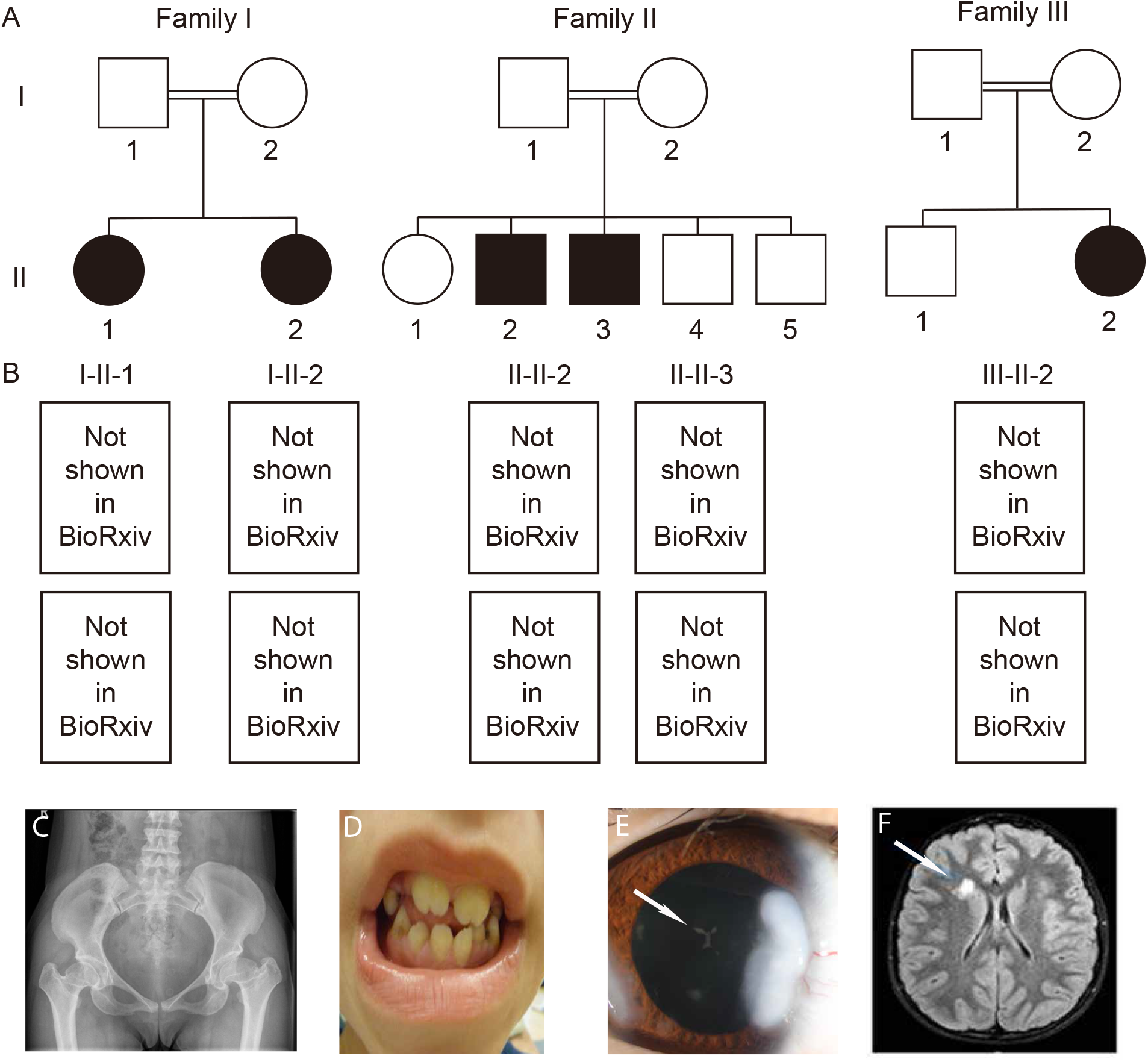
Pedigrees and pictures of the individuals studied. (A) Pedigree of three consanguineous families studied. Black boxes indicated affected individuals. Roman numerals indicating the generation are on the left and Arabic numerals indicating the individual are below each pedigree symbol. (B) Photographs of affected individuals under their corresponding pedigree symbol indicating coarse facial features including a broad nasal bridge, thick columella, and thick alae nasi. Of note, the left eye of patient II-II-2 shows phthisis bulbi of unknown etiology, as evidenced by an atrophic non-functional eye. Representative images are shown of (C) an X-ray indicating square shaped vertebral bodies and a flat pelvis, subluxation of the hips, and meta- and epiphyseal dysplasia of the femoral heads in patient III-II-2. (D) the teeth in patient II-II-3 indicating broad maxilla incisors, narrow mandible teeth, and dental enamel defects (E) the eye with a visible cataract (Cataracta polaris anterior), as indicated by a white arrow, in individual III-II-2, and (F) a brain MRI demonstrating areas of altered signal intensity as indicated by the white arrow in individual I-II-2.

In addition to the shared syndromic features described above in all three families, both affected daughters in Family I were diagnosed with congenital adrenal hyperplasia (CAH), due to 17-alpha-hydroxylase deficiency, and were found to have a homozygous familial mutation: NM_000102.3:c.286C>T; p.(Arg96Trp) in the *CYP17A1* gene (OMIM #202110) (Laflamme et al., 1996; Martin et al., 2003). The affected individuals in Families II and III do not carry mutations in *CYP17A1* or have CAH, suggesting the presence of two independent and unrelated conditions in Family I. The co-occurrence of multiple monogenic disorders is not uncommon among this highly consanguineous population (Kurolap et al., 2016).

To identify the genetic basis of this disorder, enzymatic assays related to the mucopolysaccharidosis subtypes MPS I, MPS IVA, MPS IVB, and MPSVI were tested in Families I and II and found to be normal. Enzymatic assays for mucolipidosis II/III were also normal and no pathogenic mutations were found in galactosamine-6-sulfate sulfatase (*GALNS*) in Family I. Additionally, since some of the features of patient II-II-3 were reminiscent of Noonan syndrome, Hennekam syndrome, and Aarskog-Scott syndrome, individual genes involved in these disorders were analyzed in Family II, but no pathogenic mutation was identified. In patient III-II-2, Williams-Beuren syndrome was excluded in childhood. Additionally, direct molecular testing at presentation in adulthood excluded Leri-Weill syndrome, Alstrom disease, and mutations in *FGFR3*.

Given the negative results of targeted genetic testing, WES and CNV analysis was performed for the affected individuals from all three families. Five homozygous candidate variants were identified in Family I, including the *CYP17A1* (p.Arg96Trp) mutation that is the cause of the CAH (Laflamme et al., 1996; Martin et al., 2003), but is not known to cause the other phenotypes. The remaining four variants affected the genes *ATF4*, *DNAH14*, *PLEKHA7*, and *PIK3C2A* (Table 2). In Family II, homozygous missense variants were identified in *KIAA1549L*, *METAP1*, and *PEX2*, in addition to a homozygous deletion in *PIK3C2A* that encompassed exons 1-24 out of 32 total exons (Table 2). The deletion was limited to *PIK3C2A* and did not affect the neighboring genes. Sequence analysis of Family III showed a homozygous missense variant in *PTH2R*, nonsense variant in *DPRX*, and splice site variant in *PIK3C2A* (Table 2).

**Table 2.**
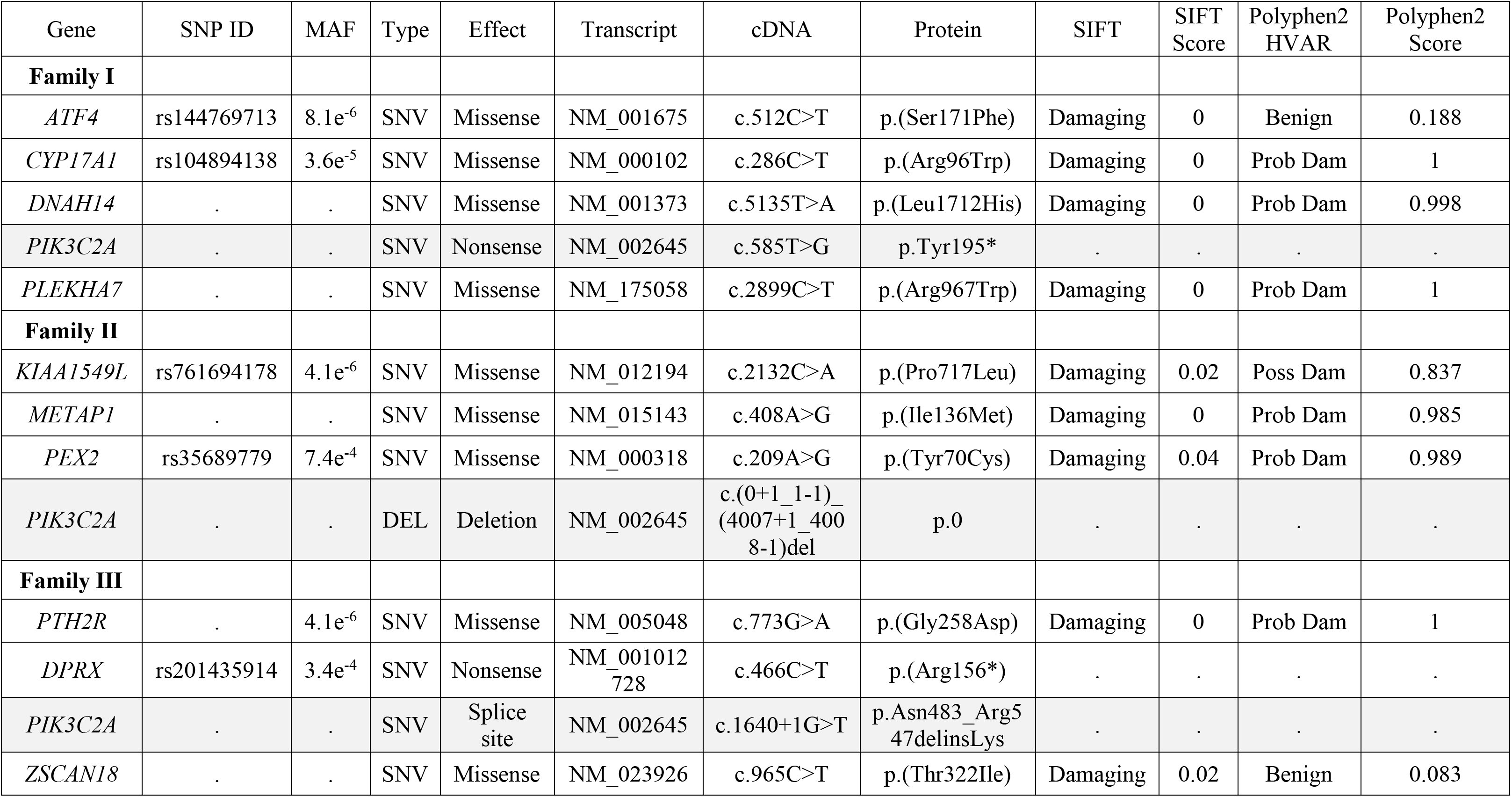
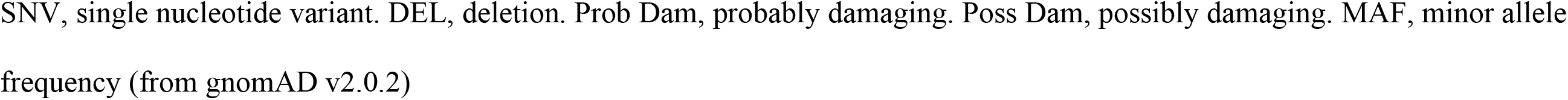
Homozygous candidate variants identified by WES.

| Gene | SNP ID | MAF | Type | Effect | Transcript | cDNA | Protein | SIFT | SIFT Score | Polyphen2 HVAR | Polyphen2 Score |
| --- | --- | --- | --- | --- | --- | --- | --- | --- | --- | --- | --- |
| **Family I** |  |  |  |  |  |  |  |  |  |  |  |
| *ATF4* | rs144769713 | 8.1e<sup>-6</sup> | SNV | Missense | NM_001675 | c.512C>T | p.(Ser171Phe) | Damaging | 0 | Benign | 0.188 |
| *CYP17A1* | rs104894138 | 3.6e<sup>-5</sup> | SNV | Missense | NM_000102 | c.286C>T | p.(Arg96Trp) | Damaging | 0 | Prob Dam | 1 |
| *DNAH14* | . | . | SNV | Missense | NM_001373 | c.5135T>A | p.(Leu1712His) | Damaging | 0 | Prob Dam | 0.998 |
| *PIK3C2A* | . | . | SNV | Nonsense | NM_002645 | c.585T>G | p.Tyr195* | . | . | . | . |
| *PLEKHA7* | . | . | SNV | Missense | NM_175058 | c.2899C>T | p.(Arg967Trp) | Damaging | 0 | Prob Dam | 1 |
| **Family II** |  |  |  |  |  |  |  |  |  |  |  |
| *KIAA1549L* | rs761694178 | 4.1e<sup>-6</sup> | SNV | Missense | NM_012194 | c.2132C>A | p.(Pro717Leu) | Damaging | 0.02 | Poss Dam | 0.837 |
| *METAP1* | . | . | SNV | Missense | NM_015143 | c.408A>G | p.(Ile136Met) | Damaging | 0 | Prob Dam | 0.985 |
| *PEX2* | rs35689779 | 7.4e<sup>-4</sup> | SNV | Missense | NM_000318 | c.209A>G | p.(Tyr70Cys) | Damaging | 0.04 | Prob Dam | 0.989 |
| *PIK3C2A* | . | . | DEL | Deletion | NM_002645 | c.(0+1_1-1)_<br>(4007+1_4008-1)del | p.0 | . | . | . | . |
| **Family III** |  |  |  |  |  |  |  |  |  |  |  |
| *PTH2R* | . | 4.1e<sup>-6</sup> | SNV | Missense | NM_005048 | c.773G>A | p.(Gly258Asp) | Damaging | 0 | Prob Dam | 1 |
| *DPRX* | rs201435914 | 3.4e<sup>-4</sup> | SNV | Nonsense | NM_001012<br>728 | c.466C>T | p.(Arg156*) | . | . | . | . |
| *PIK3C2A* | . | . | SNV | Splice site | NM_002645 | c.1640+1G>T | p.Asn483_Arg547delinsLys | . | . | . | . |
| *ZSCAN18* | . | . | SNV | Missense | NM_023926 | c.965C>T | p.(Thr322Ile) | Damaging | 0.02 | Benign | 0.083 |

SNV, single nucleotide variant. DEL, deletion. Prob Dam, probably damaging. Poss Dam, possibly damaging. MAF, minor allele frequency (from gnomAD v2.0.2)

Sequencing analyses revealed that all affected family members in the Families I, II, and III were homozygous for predicted loss-of-function variants in *PIK3C2A*, and none of the unaffected family members were homozygous for the *PIK3C2A* variants (Figure 2). The initial link between these three families with rare mutations in *PIK3C2A* was made possible through the sharing of information via the GeneMatcher website (Sobreira et al., 2015). The *PIK3C2A* deletion in Family II was confirmed by multiplex amplicon quantification. The single nucleotide *PIK3C2A* variants in Families I and III were confirmed by Sanger sequencing (Figure 2C, D).

**Figure 2.**
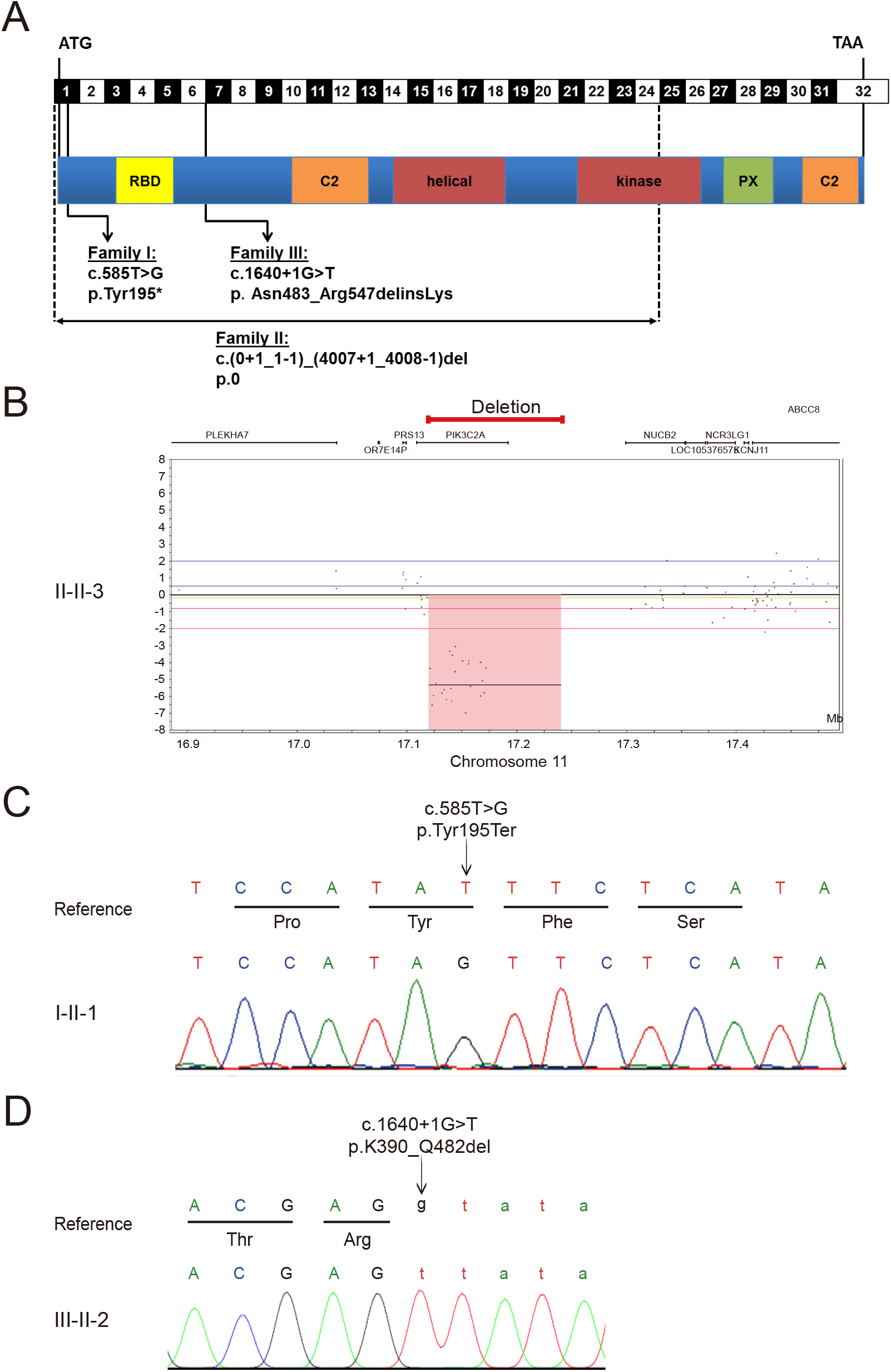
Homozygous loss-of-function mutations in *PIK3C2A*. (A) Diagram of the intron/exon and protein domain structures of *PIK3C2A* indicating the location of mutations identified in three independent consanguineous families with homozygous loss-of-function mutations in *PIK3C2A*. (B) CNV analysis confirmed a homozygous deletion encompassing exons 1-24 out of 32 total exons of *PIK3C2A*, indicated with the red line (C) Sanger sequencing confirmed homozygosity for the *PIK3C2A* c.585T variant in Family I. (D) Sanger sequencing confirmed homozygosity for the *PIK3C2A* c.1640+1 G>T variant in Family III.

In Family I, the nonsense mutation in *PIK3C2A* (p.Tyr195*) truncates 1,492 amino acids from a protein that is 1,686 amino acids. This is predicted to eliminate nearly all functional domains including the catalytic kinase domain, and is expected to trigger nonsense-mediated mRNA decay (Campa et al., 2015). Accordingly, levels of *PIK3C2A* mRNA are significantly decreased in both heterozygous and homozygous individuals carrying the p.Tyr195* variant (Figure 3A). The deletion in Family II eliminates the first 24 exons of the 32-exon *PIK3C2A* gene and is thus predicted to cause a loss of protein expression. This is consistent with a lack of *PIK3C2A* mRNA expression (Figure 3B). The variant in *PIK3C2A* in Family III affects an essential splice site (c.1640+1G>T) that leads to decreased mRNA levels (Figure 3C). Deep sequencing of the RT-PCR products revealed 4 alternative transcripts in patient-derived lymphocytes (p.[Asn483_Arg547delinsLys, Ala521Thrfs*4, Ala521_Glu568del, and Arg547SerinsTyrIleIle*]) of which the transcript encoding p. Asn483_Arg547delinsLys that skips both exons 5 and 6 was also observed in patient’s fibroblasts (Figure S2). Although this transcript remains in-frame, no PIK3C2A protein was detected by Western blotting (Figure 3D). This is consistent with Families I and II, for which Western blotting also failed to detect any full-length PIK3C2A in fibroblasts from the affected homozygous children (Figure 3E). Thus, all three *PIK3C2A* variants likely encode loss-of-function alleles. Importantly, among the 141,456 WES and whole genome sequences from control individuals in the Genome Aggregation Database (gnomAD v2.1) (Lek et al., 2016), none are homozygous for loss-of-function mutations in *PIK3C2A*, which is consistent with total PIK3C2A deficiency causing severe early onset disease.

**Figure 3.**
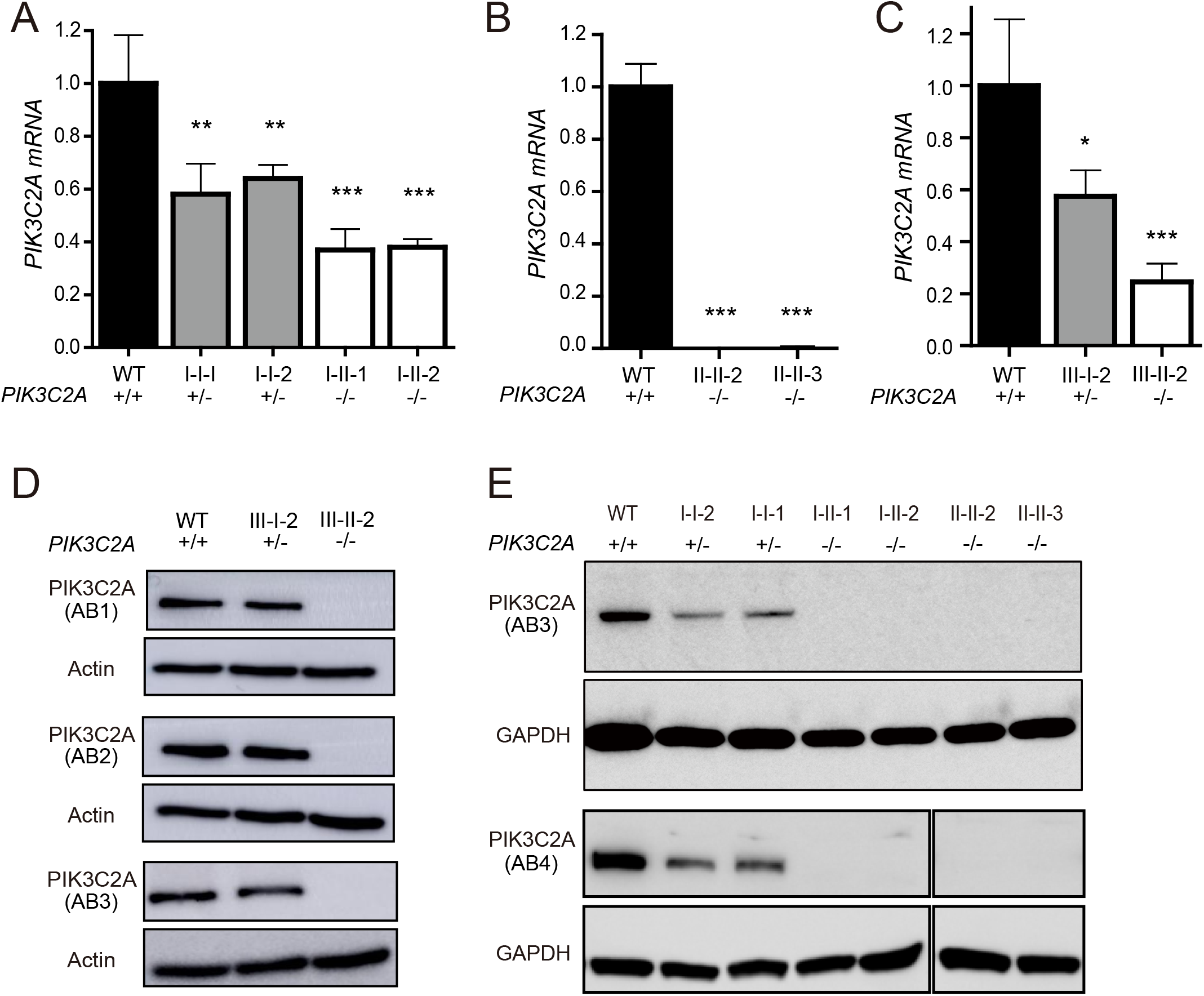
Protein and mRNA levels of PIK3C2A in patient-derived cells. *PIK3C2A* mRNA levels were detected by qRT-PCR in patient derived fibroblasts from (A) Family I, (B) Family II, and (C) Family III. (D, E) Whole cell lysates from fibroblasts of healthy controls (WT), heterozygous parents, and affected individuals from (D) Family III and (E) Families I and II were analyzed by Western blotting for PIK3C2A and the loading controls Actin or GAPDH. Immunogen of anti-PIK3C2A antibodies (AB1-AB4) are detailed in Table S4. * indicates p < 0.05. ** indicates p < 0.01. *** indicates p < 0.0001. qRT-PCR data is represented as mean ± SEM (n=3-4 technical replicates per sample).

To test whether the observed loss-of-function mutations in *PIK3C2A* cause cellular phenotypes consistent with loss of PIK3C2A function, we examined PI metabolism, cilia formation and function, and cellular proliferation rates. PIK3C2A deficiency in the patient-derived fibroblasts decreased the levels of PI(3,4)P2 throughout the cell (Figure 4A) as well as decreased the levels of PI(3)P at the ciliary base (Figures 4B, S3A). The reduction in PI(3)P at the ciliary base was associated with a reduction in ciliary length (Figure 5A), although the percentage of ciliated cells was not altered (Figure 5B). Additional cilia defects include a reduction in the levels of RAB11 at the ciliary base (Figures 5C, S3B), and increased accumulation of IFT88 along the length of the cilium (Figures 5D, S3C) that are suggestive of defective trafficking of ciliary components. Finally, the proliferative capacity of PIK3C2A deficient cells was reduced relative to control cells (Figure 6).

**Figure 4.**
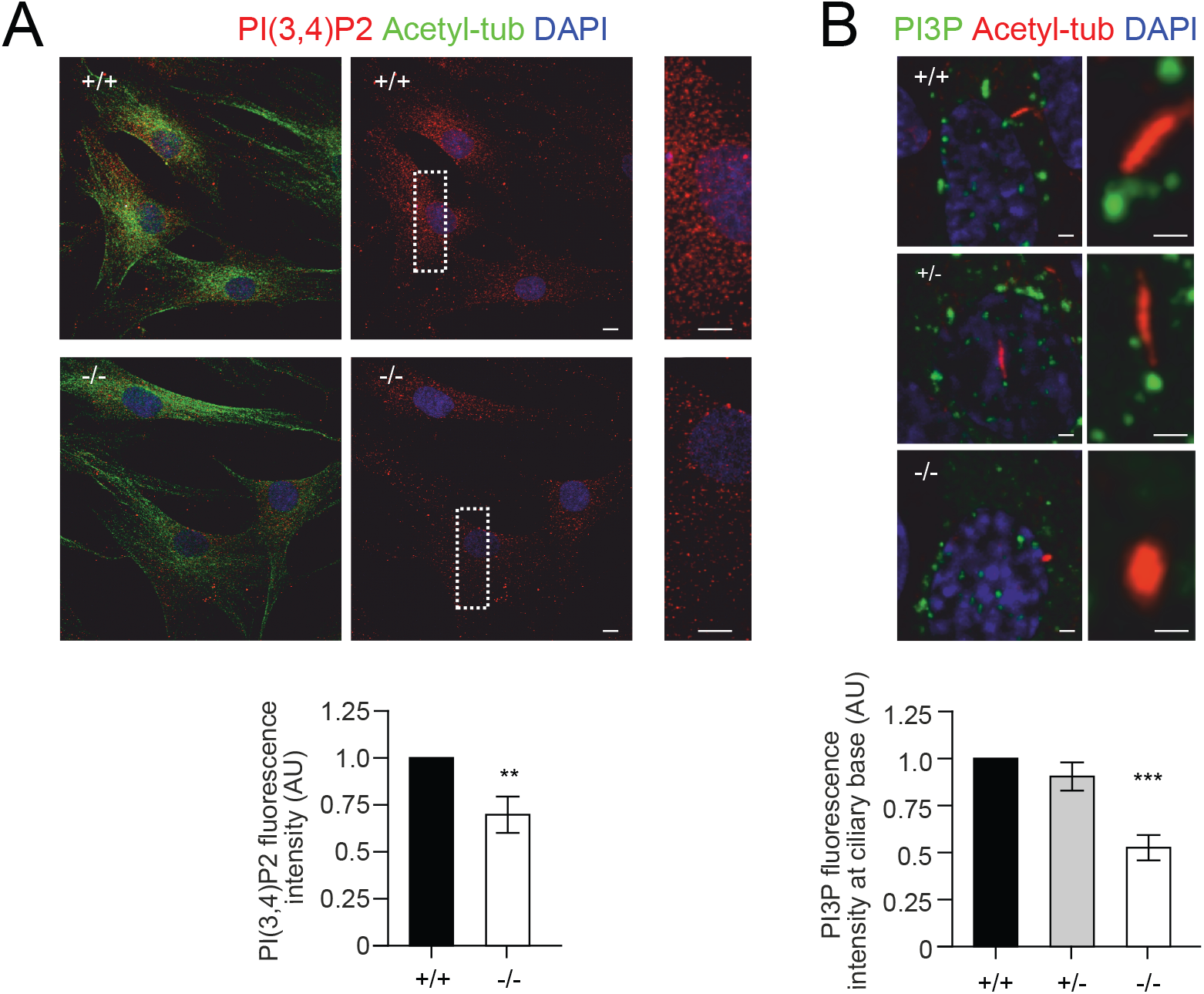
Impaired PI metabolism in patient-derived fibroblasts. Quantification of PI(3,4)P2 and PI(3)P were analysed and normalized on whole cell fluorescence. (A) Immunofluorescence analysis of cellular PI(3,4)P2 levels. Results showed that PI(3,4)P2 is significantly reduced throughout the cell in -/- cells compared with +/+ cells. (B) Immunofluorescence analysis of PI(3)P localization at the base of the primary cilium. Results showed that PI(3)P is significantly reduced at the base of the primary cilium in -/- cells compared with +/+ and or +/- cells. Nuclei are stained with DAPI. ** indicates p < 0.01. *** indicates p < 0.0001.

**Figure 5.**
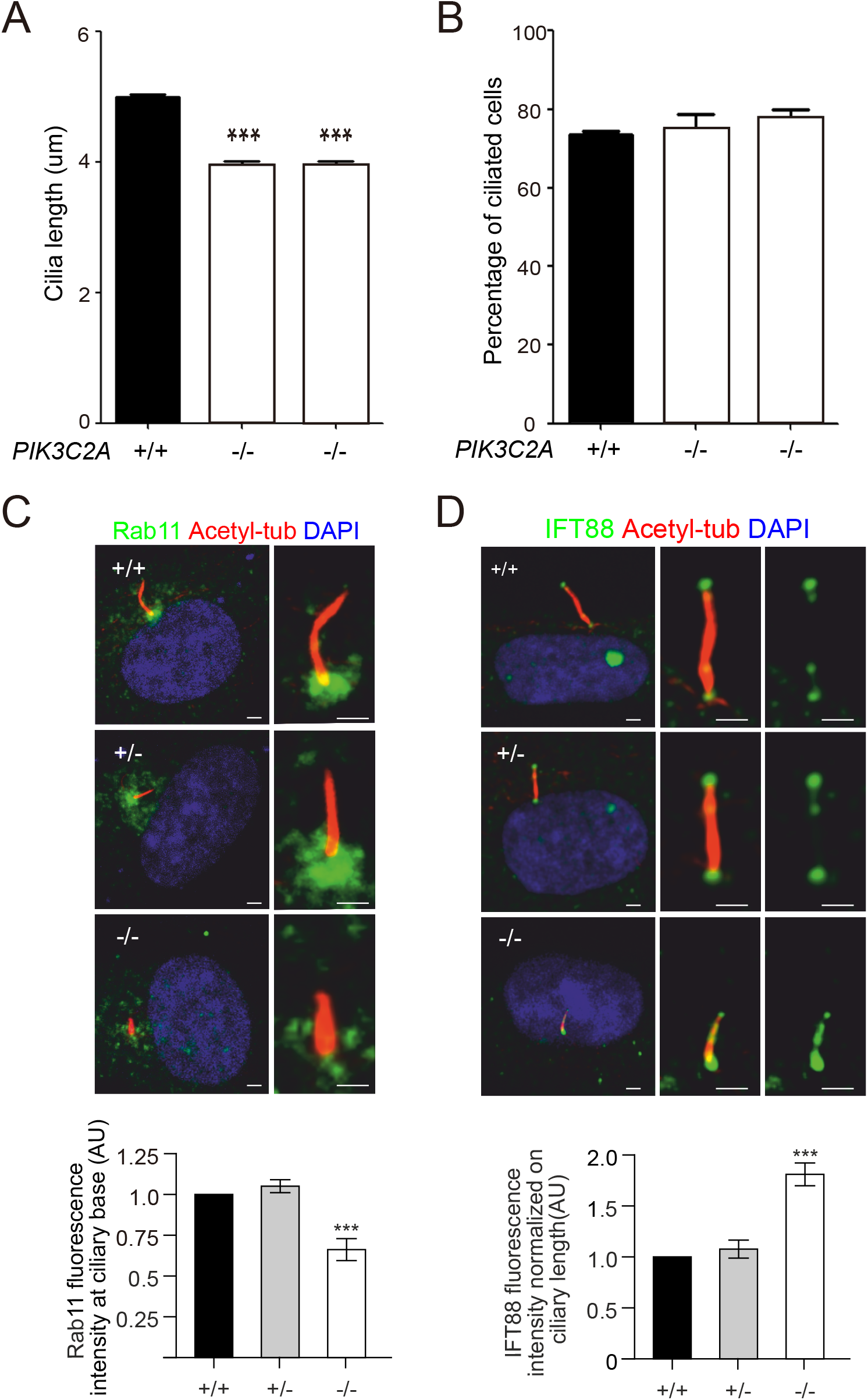
Ciliary defects due to PIK3C2A deficiency in patient-derived fibroblasts. (A) Cilia length and (B) cilia number were determined in primary fibroblasts from two affected individuals and three unrelated controls. Data is represented as mean ± SEM (n>300/sample). (C) Immunofluorescence analysis of RAB11 localization at the base of the primary cilium. Results showed that RAB11 is significantly reduced at the base of the primary cilium in -/- cells compared with +/+ and or +/- cells. (D) Immunofluorescence analysis of IFT88 localization within the primary cilia. Results showed that IFT88 is significantly increased along the primary cilium in -/- cells compared with +/+ and or +/- cells, suggesting a defective trafficking of ciliary components. Quantification of IFT88 and RAB11 were normalized on whole cell fluorescence. *** indicates p < 0.0001.

**Figure 6.**
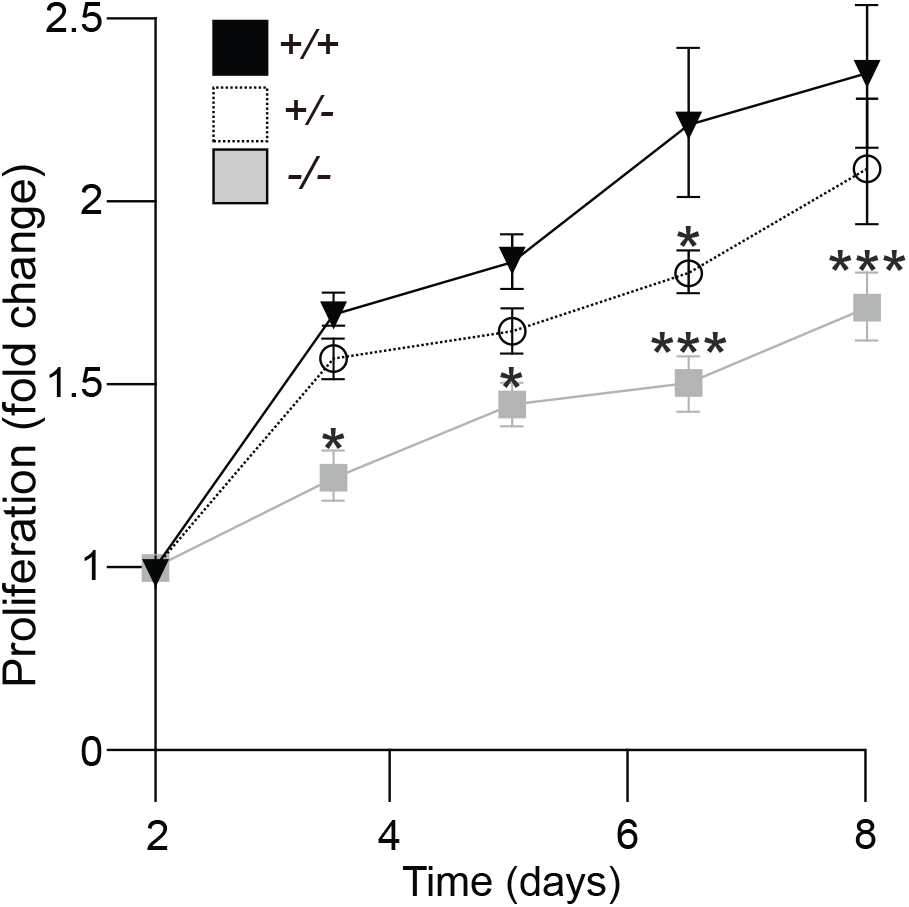
PIK3C2A deficiency causes delayed proliferation rates in patient-derived fibroblasts. Proliferation curve of primary fibroblasts isolated from *PIK3C2A* WT (+/+), heterozygous (+/-), and homozygous (-/-) individuals. Values are reported as the mean ± SEM. * indicates p < 0.05, *** indicates p < 0.0001.

As *PIK3C2A* is a member of the class II PI3K family, we tested whether the expression of the other family members *PIK3C2B* and *PIK3C2G* were altered by PIK3C2A deficiency. The expression of *PIK3C2G* was not detected by qRT-PCR in either patient-derived or control primary fibroblasts. This is consistent with the relatively restricted expression pattern of this gene in the GTEx portal (GTEx Consortium, 2013), with expression largely limited to stomach, skin, liver, esophagus, mammary tissue, and kidney, but absent in fibroblast cells and most other tissues. In contrast, *PIK3C2B* expression was detected, with both mRNA and protein levels significantly increased in PIK3C2A deficient cells (Figure 7A-C). Downregulation of *PIK3C2A* using an inducible shRNA in HeLa cells also resulted in elevated levels of PIK3C2B (Figure 7D). Together, these data are consistent with increased levels of PIK3C2B serving to partially compensate for PIK3C2A deficiency.

**Figure 7.**
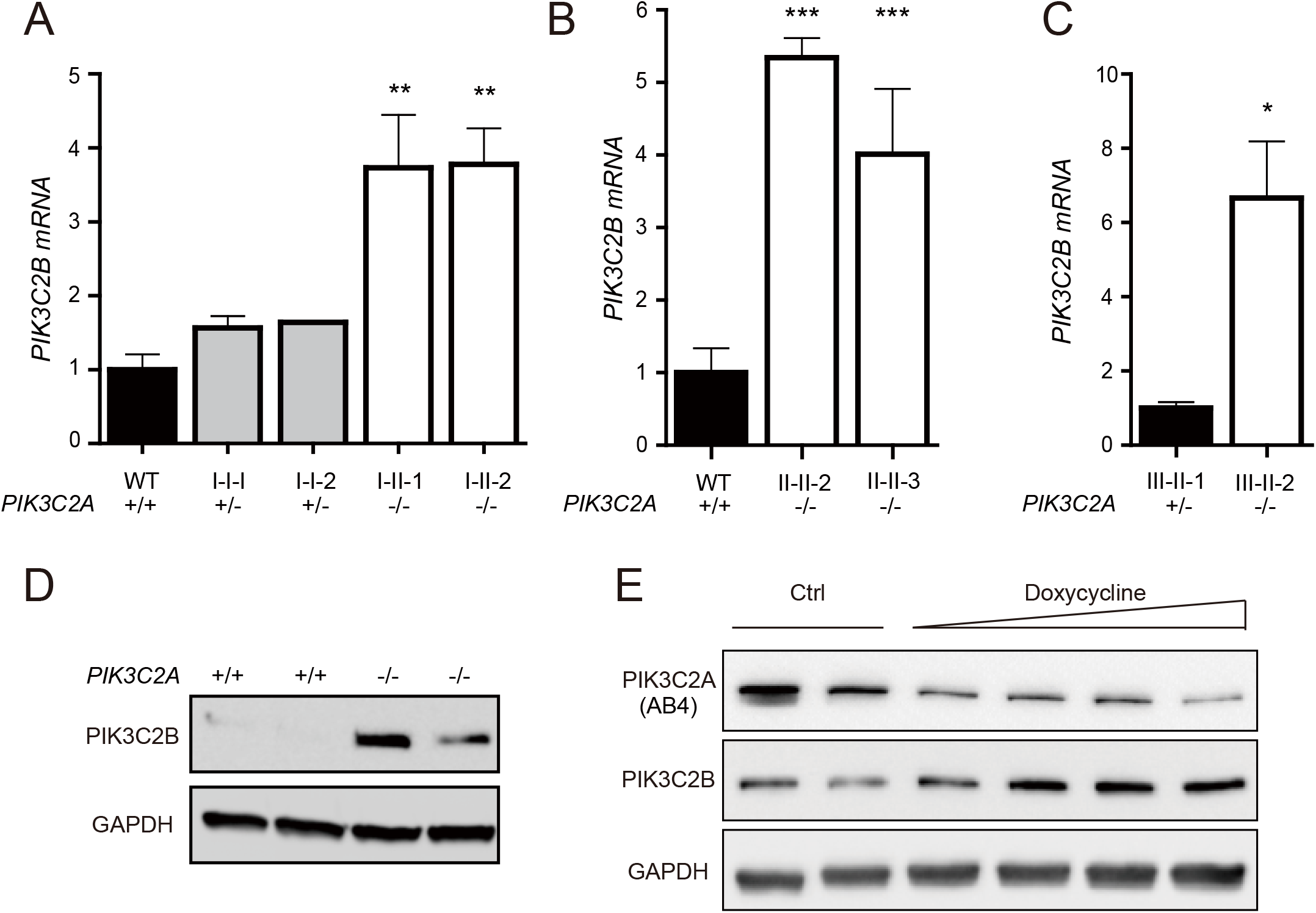
PIK3C2B levels are increased by PIK3C2A deficiency. *PIK3C2B* mRNA levels were detected by qRT-PCR in (A) Family I, (B) Family II, and (C) Family III. qRT-PCR data is represented as mean ± SEM (n=3). (D) PIK3C2B protein levels were detected by Western blotting in Family I. (E) PIK3C2A and PIK3C2B protein levels were analyzed in HeLa cells by Western blotting following doxycycline inducible shRNA mediated knockdown of PIK3C2A. * indicates p < 0.05, ** indicates p < 0.01. *** indicates p < 0.0001.

## Discussion

Here we describe the identification of three independent families with homozygous loss-of-function mutations in *PIK3C2A* resulting in a novel syndrome consisting of short stature, cataracts, secondary glaucoma, and skeletal abnormalities among other features. Patient-derived fibroblasts had decreased levels of PI(3,4)P2 and PI(3)P, shortening of the cilia and impaired ciliary protein localization, and reduced proliferation capacity. Thus, based on the loss-of-function mutations in *PIK3C2A*, the phenotypic overlap between the three independent families, and the patient-derived cellular data consistent with previous studies of PIK3C2A function, we conclude that loss-of-function mutations in *PIK3C2A* cause this novel syndrome.

The identification of *PIK3C2A* loss-of-function mutations in humans represents the first mutations identified in any class II PI-3-kinase in a disorder with a Mendelian inheritance, and thus sheds light into the biological role of this poorly understood class of PI3Ks (Jean and Kiger, 2014; Vanhaesebroeck et al., 2016). This is significant not only for understanding the role of *PIK3C2A* in rare monogenic disorders, but also the potential contribution of common variants in *PIK3C2A* in more genetically complex disorders. There are now numerous examples where severe mutations in a gene cause a rare Mendelian disorder, whereas more common variants in the same gene, with a less deleterious effect on protein function, are associated with polygenic human traits and disorders (Blair et al., 2013; Lupski et al., 2011; Marouli et al., 2017). For example, severe mutations in *PPARG* cause monogenic lipodystrophy, whereas less severe variants are associated with complex polygenic forms of lipodystrophy (Lotta et al., 2016; Semple et al., 2011). In the case of PIK3C2A deficiency, the identification of various neurological features including developmental delay, selective mutism, and the brain abnormalities detected by MRI (Table S1) may provide biological insight into the mechanisms underlying the association between common variants in *PIK3C2A* and schizophrenia (Goes et al., 2015; Ruderfer et al., 2014; Schizophrenia Psychiatric Genome-Wide Association Study (GWAS) Consortium, 2011).

Other monogenic disorders of phosphoinositide metabolism include Lowe’s syndrome and Joubert syndrome, which can be caused by mutations in the inositol polyphosphate 5-phosphatases *OCRL* and *INPP5E*, respectively (Conduit et al., 2012). All three of these disorders of PI metabolism affect some of the same organ systems, namely the brain, eye, and kidney. However, the phenotype associated with mutations in *INPP5E* is quite distinct, and includes cerebellar vermis hypodysplasia, coloboma, hypotonia, ataxia, and neonatal breathing dysregulation (Travaglini et al., 2013). In contrast, the phenotypes associated with Lowe’s syndrome share many of the same features with PIK3C2A deficiency including congenital cataracts, secondary glaucoma, kidney defects, skeletal abnormalities, developmental delay, and short stature (Bökenkamp and Ludwig, 2016; Staiano et al., 2015). The enzyme defective in Lowe’s syndrome, OCRL, is functionally similar to PIK3C2A as well, as it is also required for membrane trafficking and ciliogenesis (Mehta et al., 2014). The similarities between Lowe’s syndrome and PIK3C2A deficiency suggest that similar defects in phosphatidylinositol metabolism may underlie both disorders. In addition to Lowe’s syndrome, there is partial overlap between PIK3C2A deficiency and yet other Mendelian disorders of PI metabolism such as the early-onset cataracts in patients with INPP5K deficiency (Osborn et al., 2017; Wiessner et al., 2017), demonstrating the importance of PI metabolism in lens development.

The viability of humans with PIK3C2A deficiency is in stark contrast to mouse *Pik3c2a* knockout models that result in growth retardation by e8.5 and embryonic lethality between e10.5-11.5 due to vascular defects (Yoshioka et al., 2012). One potential explanation for this discrepancy is functional differences between human *PIK3C2A* and the mouse ortholog. However, the involvement of both human and mouse PIK3C2A in cilia formation, PI metabolism, and cellular proliferation suggests a high degree of functional conservation at the cellular level (Franco et al., 2014; Gulluni et al., 2017). An alternate possibility is that the species viability differences associated with PIK3C2A deficiency result from altered compensation from other PI metabolizing enzymes. For instance, there are species-specific differences between humans and mice in the transcription and splicing of the OCRL homolog *INPP5B* that may uniquely contribute to PI metabolism in each species (Bothwell et al., 2010). Alternately, PIK3C2B levels were significantly increased in human PIK3C2A deficient cells, including both patient-derived cells and HeLa cells surviving *PIK3C2A* deletion, suggesting that this may partially compensate for the lack of PIK3C2A in humans, although it remains to be determined whether a similar compensatory pathway exists in mice.

It is intriguing that both PIK3C2A and OCRL have important roles in primary cilia formation (Franco et al., 2014; Luo et al., 2012; Prosseda et al., 2017). Primary cilia are evolutionary conserved microtubule-derived cellular organelles that protrude from the surface of most mammalian cell types. Primary cilia formation is initiated by a cascade of processes involving the targeted trafficking and docking of Golgi-derived vesicles near the mother centriole. They play a pivotal role in a number of processes, such as left-right patterning during embryonic development, cell growth, and differentiation. Abnormal phosphatidylinositol metabolism results in ciliary dysfunction (Bielas et al., 2009), including loss of PIK3C2A that impairs ciliogenesis in mouse embryonic fibroblasts, likely due to defective trafficking of ciliary components (Franco et al., 2014). The importance of primary cilia in embryonic development and tissue homeostasis has become evident over the two past decades, as a number of proteins which localize to the cilium harbor defects causing syndromic diseases, collectively known as ciliopathies (Braun and Hildebrandt, 2017; Reiter and Leroux, 2017). Hallmark features of ciliopathies share many features with PIK3C2A deficiency and include skeletal abnormalities, progressive vision and hearing loss, mild to severe intellectual disabilities, polydactyly, and kidney phenotypes. Further work and the identification of additional patients with mutations in PIK3C2A will continue to improve our understanding of the genotype-phenotype correlation associated with PIK3C2A deficiency. However, the identification of the first patients with PIK3C2A deficiency establishes a role for PIK3C2A in neurological and skeletal development, as well as vision, and growth and implicates loss-of-function PIK3C2A mutations as a potentially new cause of a cilia-associated disease.

## Material and Methods

### Human studies

The study was approved by the ethics committees of Rambam Hospital, Haifa, Israel, and University Hospital, Erlangen, Germany and was in accordance with the regulations of the University Medical Center Groningen’s ethical committee. Informed consent was obtained from all participants.

### Whole exome sequencing

Whole exome sequencing (WES) of two patients from Family I was performed using DNA (1µg) extracted from whole blood and fragmented and enriched using the Truseq DNA PCR Free kit (Illumina). Samples were sequenced on a HiSeq2500 (Illumina) with 2×100bp read length and analyzed as described (Chen et al., 2018). Raw fastq files were mapped to the reference human genome GRCh37 using BWA (Li and Durbin, 2009) (v.0.7.12). Duplicate reads were removed by Picard (v. 1.119) and local realignment and base quality score recalibration was performed following the GATK pipeline (McKenna et al., 2010) (v. 3.3). The average read depth was 98x (I-II-1) and 117x (I-II-2). HaplotypeCaller was used to call SNPs and indels and variants were further annotated with Annovar (Wang et al., 2010). Databases used in Annovar were RefSeq (Pruitt et al., 2007), Exome Aggregation Consortium (ExAC) (Lek et al., 2016) (v. exac03), ClinVar (Landrum et al., 2016) (v. clinvar_20150330) and LJB database (Liu et al., 2011) (v. ljb26_all). Exome variants in Family I were filtered out if they were not homozygous in both affected individuals, had a population allele frequency greater than 0.1% in either the ExAC database (Lek et al., 2016) or the Greater Middle East Variome Project (Scott et al., 2016), and were not predicted to be deleterious by either SIFT (Kumar et al., 2009) or Polyphen2 (Adzhubei et al., 2010).

Whole exome sequencing was performed on the two affected individuals of Family II and both their parents essentially as previously described (Neveling et al., 2013). Target regions were enriched using the Agilent SureSelectXT Human All Exon 50Mb Kit. Whole-exome sequencing was performed on the Illumina HiSeq platform (BGI Europe) followed by data processing with BWA (Li and Durbin, 2009) (read alignment) and GATK (McKenna et al., 2010) (variant calling) software packages. Variants were annotated using an in-house developed pipeline. Prioritization of variants was done by an in-house designed ‘variant interface’ and manual curation.

The DNAs of Family III were enriched using the SureSelect Human All Exon Kit v6 (Agilent) and sequenced on an Illumina HiSeq 2500 (Illumina). Alignment, variant calling, and annotation were performed as described (Hauer et al., 2018). The average read depth was 95x (III-II-2), 119x (III-I-1) and 113x (III-I-2). Variants were selected that were covered by at least 10% of the average coverage of each exome and for which at least 5 novel alleles were detected from 2 or more callers.

All modes of inheritance were analyzed (Hauer et al., 2018). Variants were prioritized based on a population frequency of 10^-3^ or below (based on the ExAC database (Lek et al., 2016) and an inhouse variant database), on the evolutionary conservation, and on the mutation severity prediction.

All candidate variants in Families I, II, and III were confirmed by Sanger sequencing (primers listed in Table S3).

### Copy number variant (CNV) analysis

Microarray analysis for CNV detection in Family I was performed using a HumanOmni5-Quad chip (Illumina). SNP array raw data was mapped to the reference human genome GRCh37 and analyzed using GenomeStudio (v. 2011/1). Signal intensity files with Log R ratio and B-allele frequency were further analyzed with PennCNV (Wang et al., 2007) (v. 2014/5/7). In Family III the diagnostic chromosomal microarray analysis was performed with an Affymetrix CytoScan HD-Array and analyzed using Affymetrix Chromosome analysis Suite-Software, compared with the Database of Genomic Variants and 820 in house controls. All findings refer to UCSC Genome Browser on Human, February 2009 Assembly (hg19), Human Genome built 37.

CNV analysis on the WES data of Families II and III were performed using CoNIFER (Krumm et al., 2012). Variants were annotated using an in-house developed pipeline. Prioritization of variants was done by an in-house designed ‘variant interface’ and manual curation as described before (Pfundt et al., 2017). Subsequent segregation analysis of the pathogenic CNV in Family II was performed with MAQ by using a targeted primer set with primers in exons 3, 10, 20 and 24 which are located within the deletion and exons 28, 32, 34 which are located outside of the deletion (Multiplex Amplicon Quantification (MAQ); Multiplicom)).

### Cell culture

Human dermal fibroblasts were obtained from sterile skin punches cultured in DMEM (Dulbecco’s Modified Eagle’s Medium) supplemented with 10 - 20% Fetal Calf Serum, 1% Sodium Pyruvate and 1% Penicillin and streptomycin (P/S) in 5% CO2 at 37°C. Control fibroblasts were obtained from healthy age-matched volunteers. Fibroblasts from passages 4–8 were used for the experiments. To measure cell proliferation, cells were detached using trypsin and counted with an Automated Cell Counter (ThermoFisher). Cells (n=2500) were plated in triplicate in 96-well plates. Viability was measured at day 2, 4, 6 and 8. Each measurement was normalized to day 0 (measured the day after plating) and expressed as a fold increase. Viability was assessed by using CellTiter-Glo Luminescent Cell Viability Assay (Promega). Three independent experiments were performed.

### Inducible knockdown of *PIK3C2A*

HeLa cells were infected with lentiviral particles containing pLKO-TET-PI3KC2A-shRNA or pLKO-TET-scramble-shRNA in six-well plates (n=50,000 cells). After two days, the medium containing lentiviral particles was replaced with DMEM 10% FBS, 1.5µg/ml puromycin. After 7 days of selection, cells were detached and 100,000 cells were plated in six-well plates in triplicate in the presence of doxycycline (0.5, 1 and 2 µg/ml). Medium containing doxycycline was replaced every 48 hours. After 10 days of doxycycline treatment, cells were lysed and analysed by Western blot.

### cDNA and quantitative real time-PCR

Total RNA was purified from primary fibroblasts using the PureLink RNA purification kit (ThermoFisher) or RNAPure peqGOLD (Peqlab). RNA was reverse transcribed into complementary DNA with random hexamer using a high-Capacity cDNA Reverse Transcription Kit (ThermoFisher). RT-PCR from lymphocytes to detect exon-skipping in family III was performed using primers flanking exon 6. The resulting product was sequenced on an Illumina HiSeq2500 (Illumina) to detect splicing variants with high sensitivity. Gene expression was quantified by SYBR Green real-time PCR using the CFX Connect Real-Time System (BioRad). Primers used are detailed in Table S3. Expression levels were calculated using the DDCT method relative to *GADPH*.

### Western Blotting

Protein was extracted from cultured primary fibroblast cells as described (Buchner et al., 2015; Knaup et al., 2017). Extracts were quantified using the DC protein assay (BioRad) or the BCA method. Equal amounts of protein were separated by SDS-PAGE and electrotransferred onto polyvinylidene difluoride membranes (Millipore). Membranes were blocked with TBST/5% fat-free dried milk and stained with antibodies as detailed in Table S4. Secondary antibodies were goat anti-rabbit (1:5,000, ThermoFisher #31460) goat anti-mouse (1:5,000, ThermoFisher #31430), goat anti-rabbit (1:2,000, Dako #P0448), and goat anti-mouse (1:2,000, Dako #P0447).

### Immunostaining

Primary fibroblasts were grown on glass coverslips to approximately 80% -90% confluency in DMEM + 10% FCS + 1% P/S, at which time the medium was replaced with DMEM without FCS for 48 hours to induce ciliogenesis. Cells were fixed in either methanol for 10 minutes at -20°C or 4% paraformaldehyde for 10 minutes at room temperature (RT). Fixed cells were washed in PBS, and incubated with 10% normal goat serum, 1% bovine serum albumin in PBS for 1 hour at RT. If cells were fixed with paraformaldehyde, blocking solutions contained 0.5% Triton X-100. Cells were incubated with primary antibody overnight at 4°C, washed in PBS, and incubated with secondary antibody including Diamidino-2-Phenylindole (DAPI) to stain nuclei for 1 hours at RT. Coverslips were mounted on glass slides with fluoromount (Science Services) and imaged on a confocal laser scanning system with a 63x objectives (LSM 710, Carl Zeiss MicroImaging). Primary antibodies are detailed in Table S4.

### Cilia analysis

To induce ciliogenesis, cells were grown in DMEM with 0 - 0.2% FCS for 48 hours. Cells were washed in PBS, then fixed and permeabilized in ice-cold methanol for 5 minutes, followed by extensive washing with PBS. After blocking in 5% Bovine Serum Albumin, cells were incubated with primary antibodies for 1.5 hours at RT and extensively washed in PBS-T. Primary antibodies used for Centrin and ARL13B are detailed in Table S4. To wash off the primary antibody, cells were extensively washed in PBS-T. Subsequently, cells were incubated with secondary antibodies, Alexa Flour 488 (1:800, Invitrogen) and Alexa Fluor 568 (1:800, Invitrogen), for 45 min followed by washing with PBS-T. Finally, cells were shortly rinsed in ddH2O and samples were mounted using Vectashield with DAPI. Images were taken using an Axio Imager Z2 microscope with an Apotome (Zeiss) at 63x magnification. Cilia were measured manually using Fiji software taking the whole length of the cilium based on ARL13B staining. At least 300 cilia were measured per sample. Cilia lengths were pooled for 3 control cell lines and compare to 2 patient-derived samples (II-II-2 and II-II-3). Statistical significance was calculated using a Student t-test.

## Supporting information

## Acknowledgements

The authors would like to thank the Genome Aggregation Database (gnomAD) and the groups that provided exome and genome variant data to this resource. A full list of contributing groups can be found at http://gnomad.broadinstitute.org/about. This research was also supported by the Genomics Core Facility of the CWRU School of Medicine’s Genetics and Genome Sciences Department. We would also like to thank the Research Institute for Children’s Health at Case Western Reserve University and its director, Dr. Mitchell Drumm, for support and guidance. This work was supported by the NIDDK (grants DK112846 and DK099533 to D.A.B.), the Sigma Xi Scientific Research Society (grant G201603152079889 to A.C.), the DFG grants: TH 896/3-3, TH 896/3-4, TH 896/6-1, SCHU 3314/1-1, the IZKF (Interdisciplinary Centre for Clinical Research of the Universität Erlangen-Nürnberg, (FAU)) project F4, the Fondazione Italiana per la Ricerca sul Cancro (grant FIRC 19421 to F.G.), and the Johannes und Frieda Marohn-Stiftung of the FAU (WIE/2015).

## Supplemental Tables

**Table S2.**
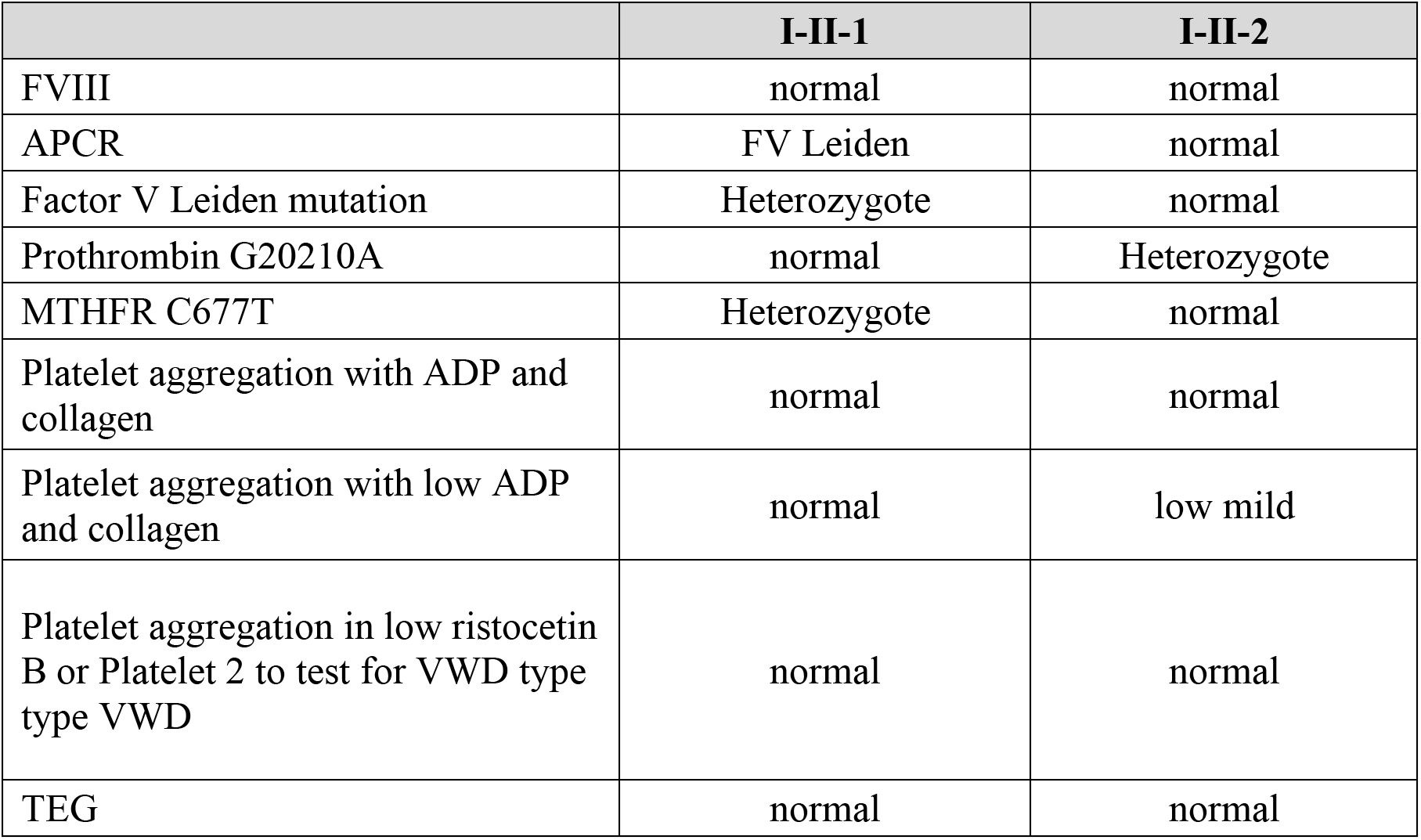
Hematological evaluation of patients in Family I.

|  | **I-II-1** | **I-II-2** |
| --- | --- | --- |
| FVIII | normal | normal |
| APCR | FV Leiden | normal |
| Factor V Leiden mutation | Heterozygote | normal |
| Prothrombin G20210A | normal | Heterozygote |
| MTHFR C677T | Heterozygote | normal |
| Platelet aggregation with ADP and collagen | normal | normal |
| Platelet aggregation with low ADP and collagen | normal | low mild |
| Platelet aggregation in low ristocetin B or Platelet 2 to test for VWD type type VWD | normal | normal |
| TEG | normal | normal |

**Table S3.**
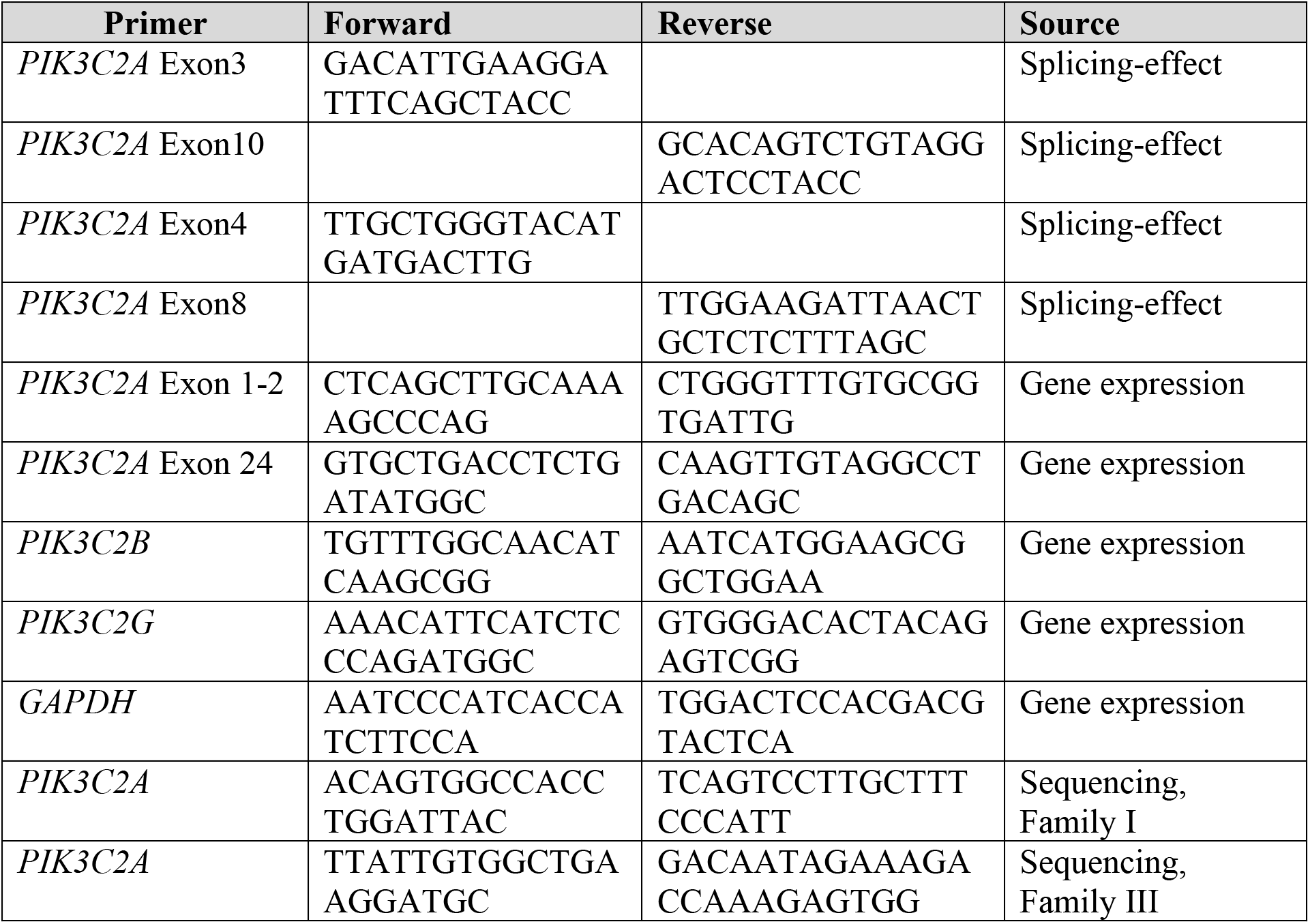
List of primers used in this study.

**Table S4.** List of antibodies used in this study.

| **Antigen** | **Host** | **Catalog #** | **Source** | **Dilution IF** | **Dilution IB** |
| --- | --- | --- | --- | --- | --- |
| PIK3C2A (AB1) | rabbit | --- | Gift from Prof. Haucke (Berlin), epitope: a.a. 2-365 | 1:200 | 1:1,000 |
| PIK3C2A (AB2) | mouse | Sc-365290 | Santa Cruz, epitope: a.a. 61-360 | 1:50 | 1:1,000 |
| PIK3C2A (AB3) | rabbit | 12402 | Cell Signaling, epitope: ~ a.a. 717 | - | 1:1,000 |
| PIK3C2A (AB4) | Rabbit | 22028-1-AP | ProteinTech, epitope: a.a. 1-338 |  | 1:1,000 |
| PIK3C2B | mouse | 611342 | BD Biosciences |  | 1:1,000 |
| Acetylated α-Tubulin (Lys40) | mouse | T7451 | Sigma | 1:300 | -- |
| Acetylated α-Tubulin (Lys40) | rabbit | 5335 | Cell Signaling | 1:200 | -- |
| IFT88 | rabbit | 13967-1-AP | ProteinTech | 1:50 | -- |
| PI(3,4)P2 | mouse | Z-P034b | Echelon | 1:150 | -- |
| PI(3)P | mouse | Z-P003 | Echelon | 1:100 | -- |
| RAB11A | rabbit | Ab65200 | Abcam | 1:100 | -- |
| Centrin | mouse | 04-1624 | Millipore | 1:500 | -- |
| ARL13B | rabbit | 17711-1-AP | Proteintech | 1:500 | -- |
| GAPDH | mouse | MA5-15738 | Thermo Fisher | -- | 1:10,000 |

IF, immunofluorescence; IB, immunoblot; a.a., amino acid

**Figure S1.**
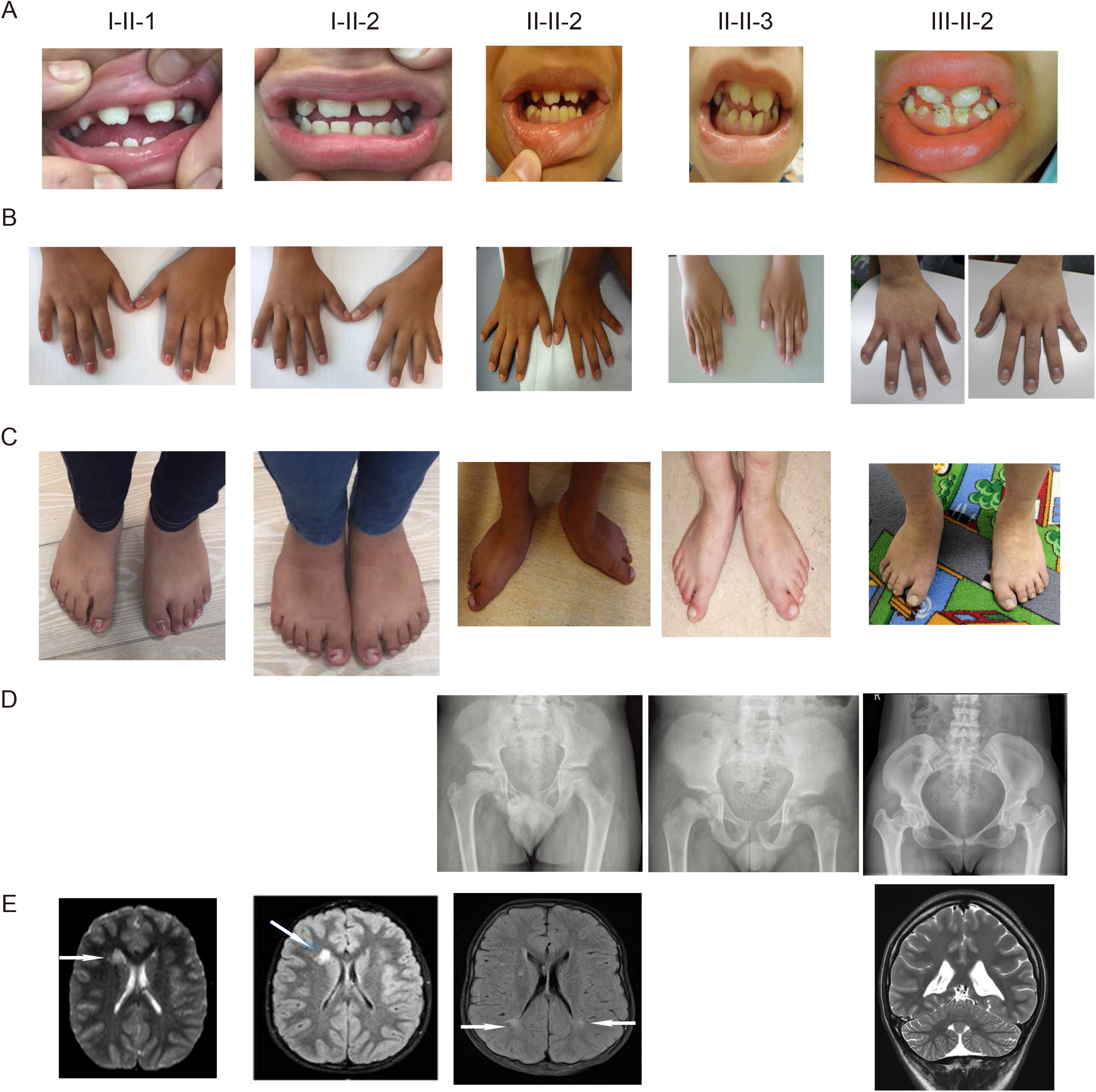
Images of individuals with PIK3C2A deficiency. Photographic images of (A) teeth, (B) hands, and (C) feet are shown from the five individuals with PIK3C2A deficiency. (D) X-Ray images of the pelvis and (E) MRI images of the brain are shown when available. White arrows in the MRI images indicate regions of altered signal intensity.

**Figure S2.**
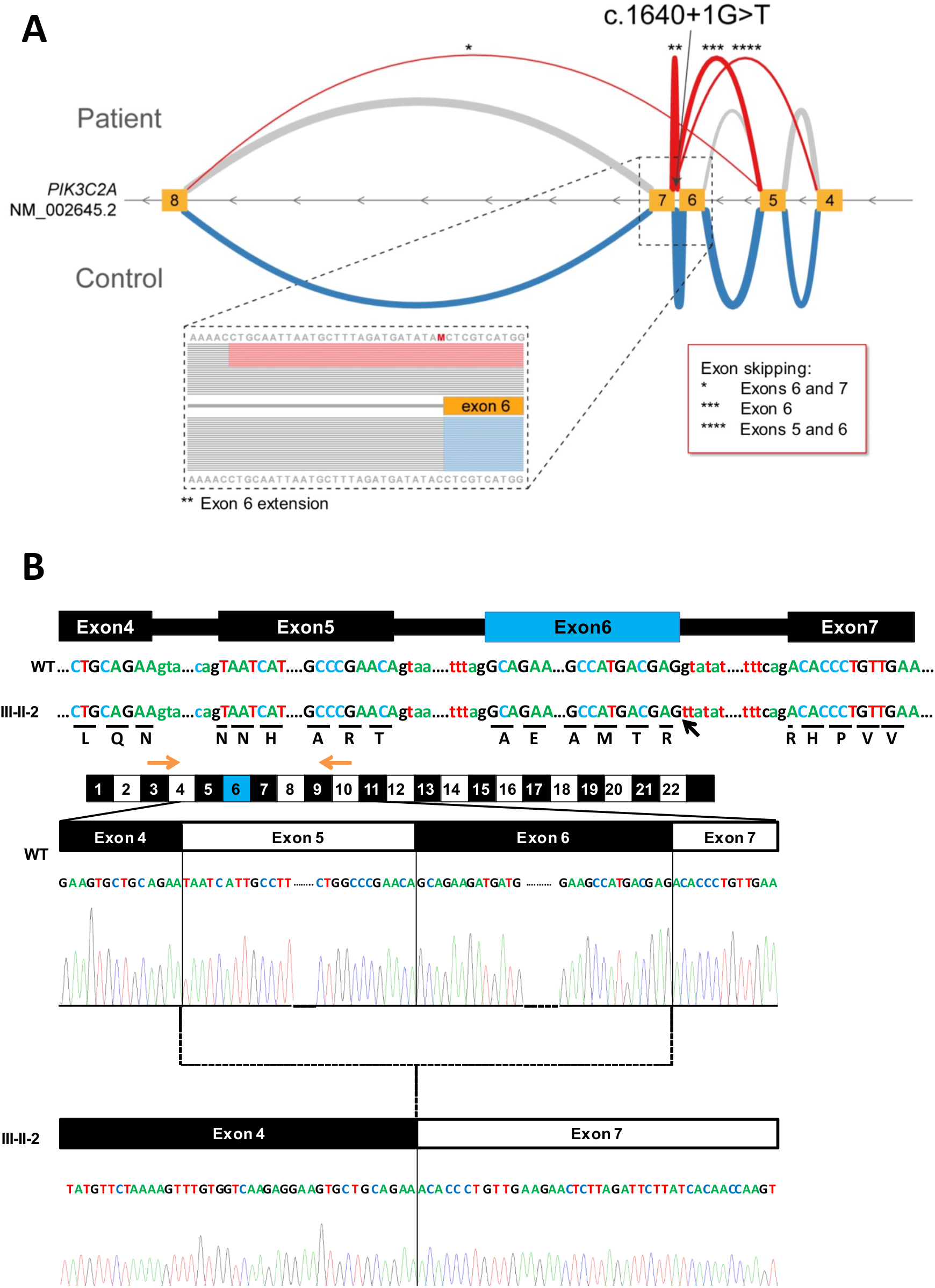
The c.1650+1G>T mutation in *PIK3C2A* disrupts the splice donor site in intron 6. (A) Deep sequencing of RT-PCR products revealed 4 alternative transcripts in lymphocytes (red lines) compared to control samples (blue lines): *: r.1561_1704del; p.Ala521_Glu568del, **: r. 1640_1641ins1640+1_1640+27; p. Arg547SerinsTyrIleIle*, ***: r.1561_1640del; p.Ala521Thrfs*4, ****: r.1449_1640del; p.Asn483_Arg547delinsLys. Exon/intron structure of *PIK3C2A* (exons: yellow boxes) with transcripts of the patient of family III compared to controls. (B) Example of sequenced RT-PCR products from cDNA of fibroblasts from wild-type control and the patient of family III using primers located in exons 3 and 10. Skipping of exons 5 and 6 is the result of the mutation. Positions of primers are indicated by orange arrows and position of the splice site mutation is indicated by a black arrow.

**Figure S3.**
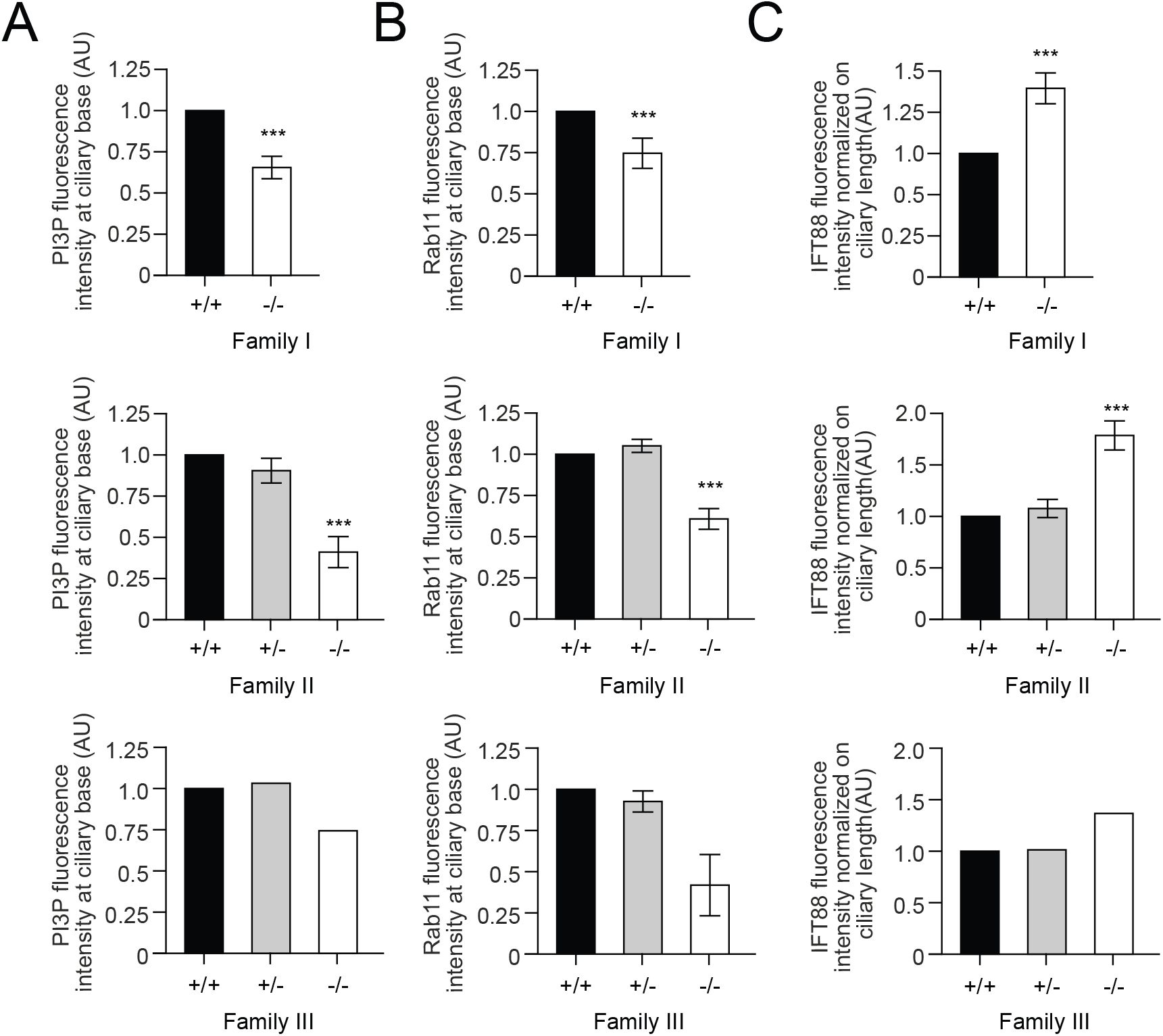
Quantification of fluorescence in patient-derived fibroblasts. Quantification of fluorescence intensity for (A) PI(3)P at the ciliary base, (B) Rab11 at the ciliary base, and (C) ciliary IFT88. Data is shown seperately for each individual family as indicated.

